# Vision shapes neural maps of space through an ancient midbrain pathway

**DOI:** 10.64898/2026.05.16.725555

**Authors:** Joshua M. Brenner, Sarah Ruediger, Cameron Wilhite, Josue M. Regalado, Yuta Senzai, Yuliya Voskobiynyk, Jeanne T. Paz, Massimo Scanziani, Riccardo Beltramo

**Affiliations:** Department of Physiology, University of California, San Francisco; San Francisco, CA, USA; Howard Hughes Medical Institute, University of California, San Francisco; San Francisco, CA, USA; Department of Molecular and Human Genetics, Baylor College of Medicine; Houston, TX, USA; Department of Neuroscience, Physiology and Pharmacology, University College London; London, UK; Department of Neuroscience, Feinberg School of Medicine, Northwestern University; Chicago, IL, USA; Gladstone Institute of Neurological Disease, Department of Neurology, University of California, San Francisco; San Francisco, CA, USA; Department of Physiology, Development and Neuroscience, University of Cambridge

## Abstract

Mammals rely on their senses to establish their position in space. Neural activity in the hippocampus maps position, yet how sensory signals reach the hippocampus remains poorly understood. Here we uncover the visual pathways informing spatial maps in the mouse hippocampus. Hippocampal activity in mice traversing a track in alternating periods of light and darkness revealed two distinct maps, one in light and one in dark. Surprisingly, distinct maps persisted following bilateral ablations of primary visual cortex, indicating that visual signals still reach the hippocampus. Conversely, blocking the ancestral pathway linking superior colliculus to lateral visual cortex markedly reduced the difference between light and dark maps. Thus, this conserved pathway relays visual information to the hippocampus, potentially explaining residual visual navigation in cortically blind humans.

## Introduction

Animals determine their position in space by synthesizing evidence from their senses, among which vision is paramount in many species. In the mammalian brain, the animal’s position in space is encoded by hippocampal place cells, whose activity is tuned to specific spatial locations known as place fields (*1–3*). Consistent with its key contribution to spatial coding, vision influences the properties of place fields: Adjusting the position of salient visual landmarks or the rate of visual flow (*4–6*), decreasing the overall salience of the visual environment (*7*), and even switching off ambient illumination can profoundly change the activity of place cells (*1*, *8*). Despite the influence of vision on hippocampal activity, the precise pathways that convey visual information to the hippocampus are still unknown.

The retina gives rise to two main image-forming streams. One pathway, prominent in mammals, reaches the primary visual cortex (V1) via the dorsolateral geniculate nucleus of the thalamus, before branching out to a suite of higher visual cortical areas including the lateral visual cortex (LVC; *9–12*). This pathway to the cortex has long been presumed to be the main conduit of visual information to the hippocampus (*9*, *13*). The other pathway proceeds through the superior colliculus (SC), a midbrain structure conserved among vertebrates, and from there reaches LVC via the pulvinar nucleus of the thalamus (*14–17*). While this pathway to the cortex has received much less attention, recent work has shown that the SC neurons that convey visual information through this pathway are widefield cells, a cell type uniquely sensitive to moving stimuli and to visual flow (*17–19*). Accordingly, LVC strongly responds to moving stimuli and visual flow, and is thus, in principle, well suited to extract information related to the animal’s self-motion (*12*, *16*, *18*). Is LVC a gateway through which visual signals shape hippocampal representations of space? If so, does this information primarily arise from the canonical geniculate-V1 pathway or from the evolutionarily older SC pathway? (Fig. 1A)

**Fig. 1.**
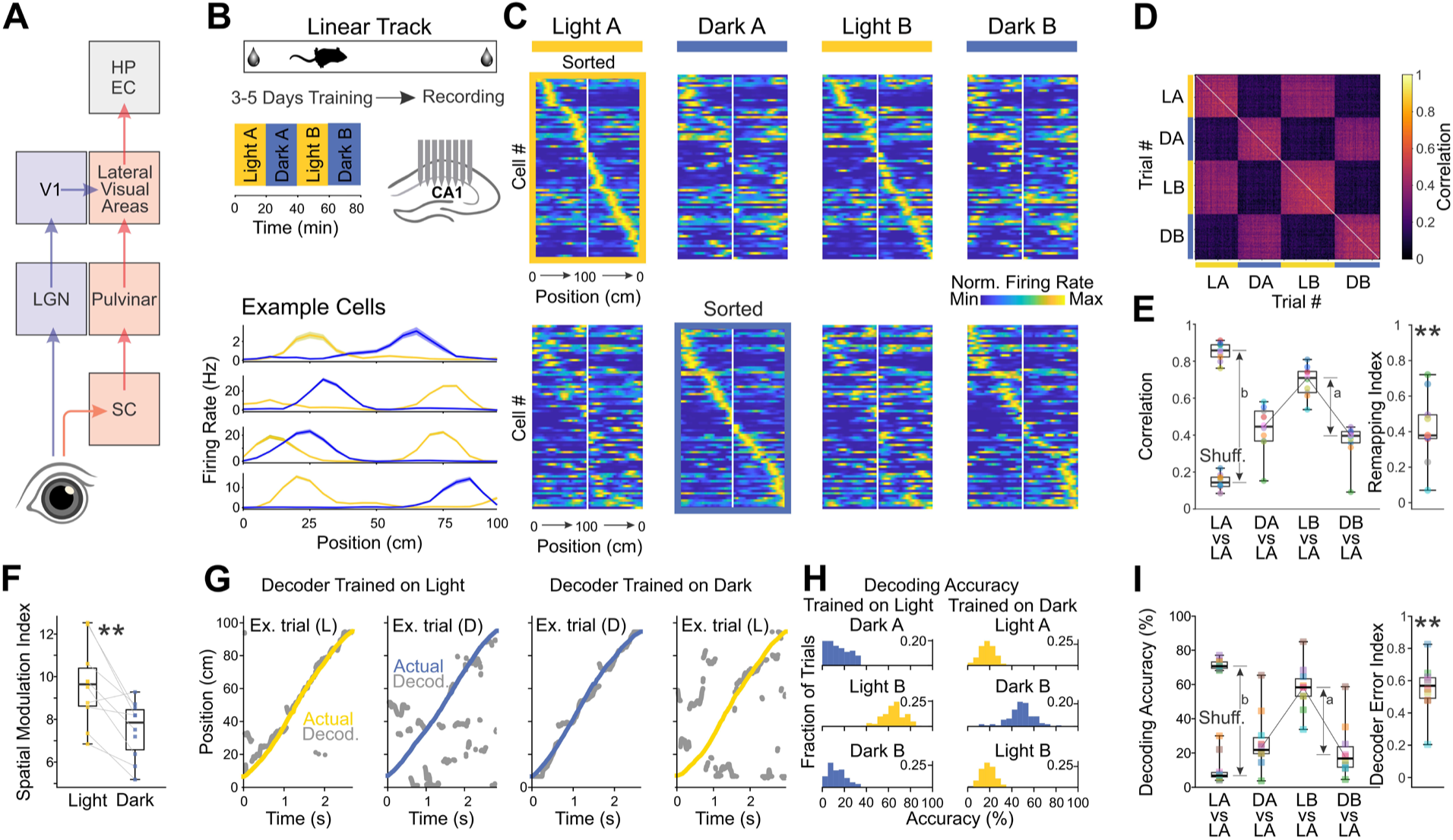
Spatial remapping in CA1 between light and darkness. **(A)** Schematic of visual pathways from retina to entorhinal cortex (EC) and hippocampus (HP) via geniculate-V1 (LGN-V1) and superior colliculus (SC) routes. **(B)** Experimental paradigm. Mice were trained for 3-5 days on a 1 m linear track with rewards at both ends. Recordings in CA1 were performed during alternating 20-min blocks of ambient light and near-complete darkness. **(C)** Normalized firing rate maps of CA1 place cells from a representative mouse, sorted by peak firing position in Light A (top) or Dark A (bottom). Maps represent concatenated leftward and rightward runs. **(D)** Pairwise Spearman correlation matrix of population activity across all trials for a representative mouse (chronologically ordered). LA/LB: Light blocks A and B; DA/DB: Dark blocks A and B. **(E)** Left: Spearman correlation of population vectors between odd and even trials within and across blocks for all mice. “Shuff.” (Shuffle) represents correlations between population vectors in odd trials and spatially randomized population vectors in even trials. Box plots indicate median (center line), interquartile range (box edges), and range (whiskers). a: difference between median correlation in same-condition (LA vs LB) and cross-condition (LA vs Average of DA and DB); b: difference between same-condition and chance (Shuffle). Right: Remapping Index, defined as the ratio a/b. N = 10 mice; **, p = 0.002 (one-sample Wilcoxon signed-rank test). **(F)** Spatial Modulation Index (SMI) of place fields in light and dark blocks for all mice. Box plots formatted as in (E). N = 10 mice; *, p = 0.006 (Wilcoxon signed-rank test). **(G)** Bayesian decoding of position. Plots show actual position (colored lines) versus decoded position (grey points) using a decoder trained on Light A (left columns) or Dark A (right columns). **(H)** Distribution of decoding accuracy for all trials in a representative mouse. **(I)** Left: Decoding accuracy across all trials for each mouse. Shuffle was calculated by randomly permuting decoded positions across spatial bins and calculating the percentage of bins in which the error fell within the actual Light A threshold. a: difference in decoding accuracy between same-condition (LA on LB) and cross-condition (LA on Dark); b: difference between same-condition and chance (Shuffle). Right: Decoder Error Index, defined as a/b. N = 10 mice; **, p = 0.002 (one-sample Wilcoxon signed-rank test).

## Results

We first established a simple and reproducible behavioral protocol that allows us to determine the impact of candidate visual pathways on hippocampal space representation (Fig. 1, A and B). To that end, we compared hippocampal maps of mice running along the same trajectory under ambient illumination and in near total darkness, that is, less than 100 isomerizations per rod per second (see methods). To minimize behavioral variability between light and dark conditions, we constrained the trajectories of the mice by training them to traverse a narrow linear track (100 × 5 cm) with a soymilk reward at each end. Furthermore, we subdivided the floor of the track into sections with distinct tactile textures to provide the animal with positional cues that do not require vision. We trained mice on this linear track in alternating twenty-minute light and dark blocks for 3–5 days until they could complete at least 200 laps in two hours. We refer to consecutive, alternating light and dark blocks as Light A, Dark A, Light B, and Dark B. This apparatus and training protocol resulted in a stereotyped locomotor behavior where running trajectory, speed, and acceleration were unchanged between light and darkness. After training, we recorded from hippocampal CA1 neurons to isolate place cells (see methods). The spatial representation of the track was markedly different between light and dark blocks. While place fields tiled the track in both conditions, switching from Light A to Dark A reconfigured the sequence of peak activation of place cells along the track. This spatial remapping was reversible—returning to ambient illumination (Light B) restored the activation sequence observed in Light A, and switching back to darkness (Dark B) changed the sequence again to match that of Dark A.

To compare the spatial activity pattern of CA1 cells between light and dark trials, we used two independent procedures: a correlation analysis (Fig. 1C) to quantify the similarity of population activity patterns across trials, and a decoding approach (*20*, *21*), in which a decoder trained on a subset of trials estimates the position of the animals on the untrained set (see methods). The stark checkerboard pattern reveals that the sequence of CA1 activation was highly correlated across light trials – irrespective of the block – but weakly correlated to dark trials, and vice versa. (Fig. 1D). Accordingly, the correlation between Light A and Dark A or Dark B blocks was smaller than that between Light A and Light B blocks (Fig. 1E; Spearman Correlation for Light A Even to Light A Odd: Median = 0.8582, IQR = 0.0660; for Light A to Dark A: Median = 0.4456, IQR = 0.1620; for Light A to Light B: Median = 0.6914, IQR = 0.114). We used this difference in correlation to compute the “remapping index” where 0 and 1 indicate no or maximal spatial remapping, respectively (see methods). The remapping index had a median value of 0.3789, with some animals showing almost maximal remapping while others only a relatively small shift (Remapping Index, Controls: Median = 0.3789, IQR = 0.1337, N = 10, p = 0.0019). Irrespective of the absolute value of the remapping index, however, in the vast majority of animals (9 out of 10) the correlation between Light A and Light B was larger than between Light A and any dark block. There was little difference detected in mouse locomotor behavior between light and dark (Fig. S1, A to E; Velocity (cm/s): Light median = 27.40, IQR = 5.11; Dark median = 27.08, IQR = 5.38; N = 10; p = 1.00. Acceleration (magnitude; cm/s^2^): Light median = 29.46, IQR = 5.21; Dark median = 30.22, IQR = 5.90; N = 10; p = 0.63). Nevertheless, to confirm that the remapping was not due to subtle variations in behavior, we specifically selected a subset of trials from light and dark such that their mean trajectories, velocities, and accelerations were nearly identical. The spatial remapping of the sequence of place fields between light and dark of these select trajectories was indistinguishable from the remapping occurring between the entire light and dark blocks (Fig. S1, F and G; All Trials, Example Mouse: Median = 0.5376, IQR = 0.0756, n = 48; Subset: Median = 0.5140, IQR = 0.0862; N = 7; p = 0.63). Thus, despite nearly identical locomotor trajectories and behaviors, CA1 formed distinct representations of the same environment in light and darkness.

To confirm the remapping between light and dark, we next used a decoder trained to predict the animal’s position on the track. A decoder trained on a subset of dark trials was equally precise at predicting the position of the animal on untrained dark trials as a decoder trained on a subset of light trials at predicting the animal’s position on untrained light trials. This, despite the fact that the spatial modulation index (SMI) of CA1 cells was slightly lower in dark (Fig. 1F and Fig. S2A; Light Median SMI = 9.64, IQR = 1.7741; Dark Median SMI = 7.85, IQR = 1.8740; N = 10; p = 0.006). Crucially, however, and consistent with the remapping between light and dark described above, the decoder trained on light trials was unable to accurately predict the animal’s position in dark trials, and vice versa (Fig. 1, G and H). We quantified the performance of a decoder trained in light to decode the position of the animal in dark through an index of decoding error, the “error index,” where a value of 0 indicates that the decoder accuracy is as good in dark as in light and a value of 1 indicates that the decoder performs at chance in dark. The decoder error index had a median of 0.568 (Fig. 1I; Decoder Error Index Control: Median = 0.568, IQR: 0.127, N = 11, p = 0.002). Furthermore, across animals, there was a tight correlation between the decoder error index and the remapping index, consistent with the idea that large remapping of place cell sequences between light and dark leads to poorer estimates of the animal’s position in dark for decoders trained in light (Pearson r: 0.896, p = 8.186e-13). These results indicate that these mice acquire two stable but distinct hippocampal representations of the linear track, one under ambient illumination in the presence of vision and one in darkness in the absence of vision. This paradigm thus allows us to test candidate pathways for visual input to reach the hippocampus.

If LVC is a conduit of visual information to the hippocampus, its ablation should impact the spatial remapping of place fields between light and darkness. Specifically, if ablation of LVC impairs the flow of visual information to the hippocampus, then the spatial representation should change little in the presence or absence of a visual input. We thus performed bilateral aspiration lesions of LVC in naive animals, trained them on the linear track as for control animals ten days after surgery, and tested the impact of light and dark on the spatial representation of the track in the hippocampus (Fig. 2, A and B). The lesions of LVC included the postrhinal cortex (POR), the lateromedial area (LM), and the laterointermediate area (LI) (*22*); the precise extent of each lesion was quantified histologically post hoc using two independent approaches (Fig. S3; LVC Profilometry: Median = 0.559 mm^3^, IQR = 0.205 mm^3^, N = 5. LVC Histology: Median = 0.550 mm^3^, IQR = 0.223 mm^3^, N = 9). Lesioned animals learned the task and performed as many trials as control animals. Furthermore, as in control animals, laterally lesioned mice formed hippocampal spatial representations that tiled the track. In contrast to control mice, however, in laterally lesioned mice changing from light to dark led to significantly less remapping. The remapping index dropped from 0.379 in control to 0.202 in lateral lesioned animals (Fig. 2, C to F; Remapping Index, Lateral Lesions: Median = 0.202, IQR = 0.112, N = 8, p = 0.016, Controls vs. Lateral lesions). Furthermore, in contrast to control animals, shifting from light to dark did not affect the average SMI of place cells (Fig. S2B; Light Median SMI = 10.10, IQR = 4.786; Dark Median SMI = 10.14, IQR = 3.823; N = 8; p = 0.461). Finally, a decoder trained on light trials and applied to dark trials performed significantly better for lesioned animals than for control mice (Fig. 2, G to I; Decoder Error Index Lateral Lesions: Median = 0.268, IQR: 0.237, N = 8, p = 0.008, Controls vs. Lateral lesions). Thus, LVC is necessary for the remapping of CA1 place fields between light and dark, demonstrating that it conveys, at least in part, the visual information that shapes spatial representation in the hippocampus.

**Fig. 2.**
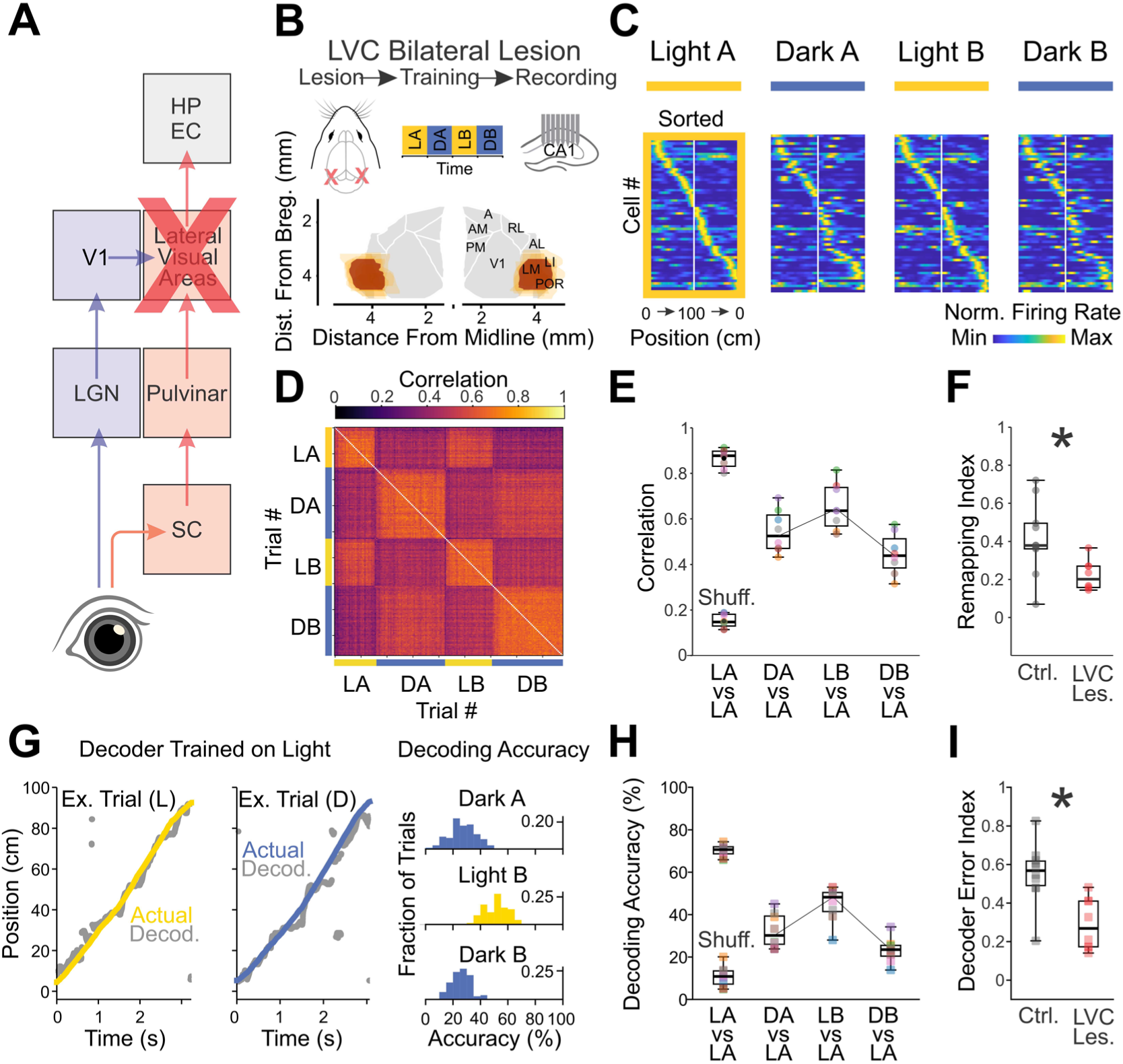
Lateral visual cortex (LVC) lesions consolidate light and dark spatial maps. **(A)** Schematic of visual pathways with bilateral lesions of the lateral visual areas (LVC) indicated by the red X. **(B)** Experimental paradigm and histology. Top: Timeline with LVC lesion surgery prior to training and recording. Bottom: Reconstruction from histology of bilateral LVC lesions superimposed on a reference map of the visual cortex generated from the Allen Common Coordinate Framework. Dark red indicates the median lesion extent across all mice (N = 8 mice), overlaid on individual lesion extents (lighter orange). **(C)** Normalized firing rate maps of CA1 place cells from a representative LVC-lesioned mouse, sorted by peak firing position in Light A. Maps represent concatenated leftward and rightward runs. **(D)** Pairwise Spearman correlation matrix of population activity across all trials for a representative LVC-lesioned mouse (chronologically ordered). **(E)** Spearman correlation of population vectors between odd and even trials within and across blocks for all LVC-lesioned mice. “Shuff.” (Shuffle) represents correlations between population vectors in odd trials and spatially randomized population vectors in even trials. Box plots indicate median (center line), interquartile range (box edges), and range (whiskers). **(F)** Remapping Index comparing Control (Ctrl., data from Figure 1) and LVC-lesioned (LVC les.) mice. Index is calculated as in Figure 1, with lower values indicating reduced remapping. N = 8 LVC mice; N = 10 control mice; *, p = 0.016 (Wilcoxon rank-sum test). **(G)** Left: Bayesian decoding of position in an LVC-lesioned mouse using a decoder trained on Light A. Plots show actual position (colored lines) versus decoded position (grey points) for a light and dark example trial. Right: Distribution of decoding accuracy for Dark A, Light B, and Dark B trials using the Light A decoder. **(H)** Decoding accuracy across all trials for each LVC-lesioned mouse (N = 8 mice). Shuffle was calculated by randomly permuting decoded positions across spatial bins and calculating the percentage of bins in which the error fell within the actual Light A threshold. **(I)** Decoder Error Index comparing Control and LVC-lesioned mice, as calculated in Figure 1. N = 8 LVC-lesioned mice; N = 10 control mice; **, p = 0.003 (Wilcoxon rank-sum test).

LVC sits at the intersection of feed-forward visual inputs from the primary visual cortex (V1) and the superior colliculus (SC) (*10*, *11*, *16*, *18*). Thus, both structures could, in principle, provide visual information used for spatial representation in the hippocampus. To test their contribution, we either ablated V1 via aspiration or blocked SC’s visual output. V1 was ablated bilaterally and, ten days later, the animals were trained on the linear track to assess the impact of light and dark on hippocampal representations, as in LVC-lesioned mice (Fig. 3A and B). The extent of V1 lesions was quantified *post hoc* using two independent approaches (Fig. S3; V1 Profilometry: Median = 0.830 mm^3^, IQR = 0.331 mm^3^, N = 5. V1 Histology: Median = 0.963 mm^3^, IQR = 0.436 mm^3^, N = 9). In contrast to LVC lesions, bilateral V1 lesions had little effect on the remapping. Furthermore, there was no significant correlation between the extent of the V1 lesions and the degree of remapping observed in each mouse (Avg. volume (profilometry) vs. remapping index: N = 5, Pearson r = −0.165, p value: 0.790). Similar to control animals, V1-lesioned mice formed hippocampal spatial representations that tiled the linear track and switching from light to dark led to a reversible remapping (Fig. 3, C to F, and Fig. S2C; Remapping Index, V1 Lesions: Median = 0.3886, IQR = 0.4498, N = 9, p = 0.7802, Controls vs. V1 lesions), albeit with no significant change in SMI between conditions (Light Median SMI = 8.03, IQR = 6.0721; Dark Median SMI = 7.87, IQR = 4.9045; N = 9; p = 0.570). Furthermore, a decoder trained on a subset of light trials was significantly more accurate in decoding the animal’s position on untrained light than dark trials, and vice versa (Fig. 3, G to I; Decoder Error Index V1: Median = 0.435, IQR = 0.306, N = 9, p = 0.653, Controls vs. V1 lesions). Thus, the visual signal that shapes spatial representations in the hippocampus remains available following V1 lesions.

**Fig. 3.**
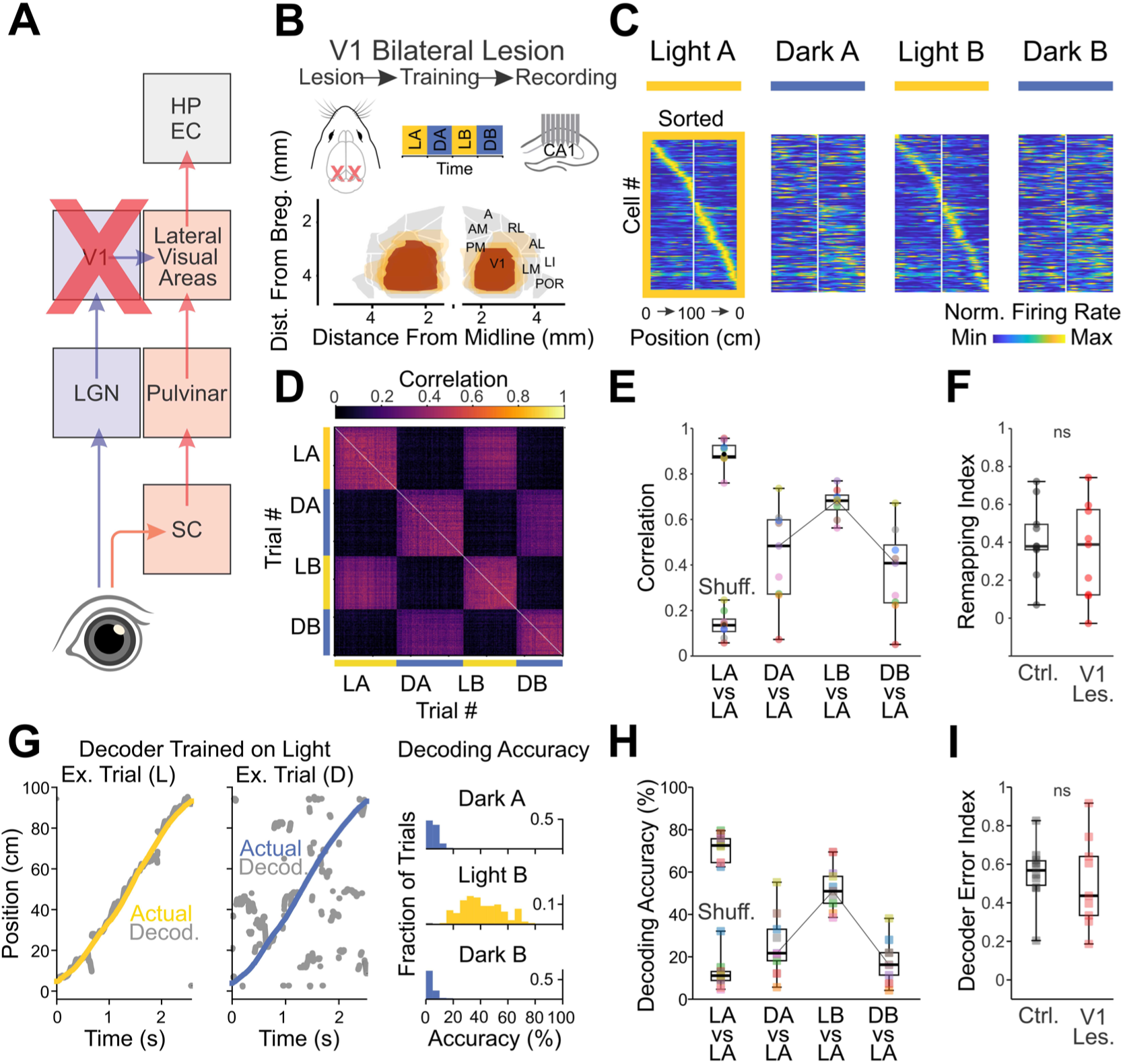
V1 lesions have no effect on separation of light and dark spatial maps. **(A)** Schematic of visual pathways with bilateral lesions of the primary visual cortex (V1) indicated by the red X. **(B)** Experimental paradigm and histology. Top: Timeline with V1 lesion surgery prior to training and recording. Bottom: Reconstruction from histology of bilateral V1 lesions superimposed on a reference map of visual cortex generated from the Allen Common Coordinate Framework. Dark red indicates the median lesion extent across all mice (N = 9 mice), overlaid on individual lesion extents (lighter orange). **(C)** Normalized firing rate maps of CA1 place cells from a representative V1-lesioned mouse, sorted by peak firing position in Light A. Maps represent concatenated leftward and rightward runs. **(D)** Pairwise Spearman correlation matrix of population activity across all trials for a representative V1-lesioned mouse (chronologically ordered). **(E)** Spearman correlation of population vectors between odd and even trials within and across blocks for all V1-lesioned mice. “Shuff.” (Shuffle) represents correlations between population vectors in odd trials and spatially randomized population vectors in even trials. Box plots indicate median (center line), interquartile range (box edges), and range (whiskers). **(F)** Remapping Index comparing Control (Ctrl., data from Figure 1) and V1-lesioned (V1 les.) mice. Index is calculated as in Figure 1, with lower values indicating reduced remapping. N = 9 V1-lesioned mice; N = 10 control mice; p = 0.780 (Wilcoxon rank-sum test). ns, not significant. **(G)** Left: Bayesian decoding of position in a V1-lesioned mouse using a decoder trained on Light A. Plots show actual position (colored lines) versus decoded position (grey points) for a light and dark example trial. Right: Distribution of decoding accuracy for Dark A, Light B, and Dark B trials using the Light A decoder. **(H)** Decoding accuracy across all trials for each V1-lesioned mouse (N = 9 mice). Shuffle was calculated by randomly permuting decoded positions across spatial bins and calculating the percentage of bins in which the error fell within the actual Light A threshold. **(I)** Decoder Error Index comparing Control and V1-lesioned mice, as calculated in Figure 1. N = 9 V1-lesioned mice; N = 10 control mice; p = 0.653 (Wilcoxon rank-sum test).

The SC relays visual signals to LVC via widefield cells, neurons located in the superficial layers of the SC that target the caudal pulvinar, the main thalamic input to LVC (*16–18*). Do widefield cells transmit visual information that contributes to the hippocampal representation of space along the track? Widefield cells are known to respond selectively to moving stimuli, yet whether they respond to static environmental stimuli that move across the visual field through the locomotion of the animal is not known (*18*, *19*). To address this question, we conditionally expressed a genetically encoded calcium indicator in widefield cells and monitored neuropil activity on the surface of the SC using a miniaturized miniscope affixed to the head of the animal (*23*; Fig. 4A). A static visual stimulus (1.2 × 1.2cm black square on white background) was presented on one side of the track while the other side was covered by a white board, for comparison (Fig. 4B; see Methods). The presence of the stimulus triggered a clear wave of activity that progressed along the anterior-posterior axis of the contralateral SC as the mice ran past the stimulus (Fig. 4C; Movie S1). The response was absent when the animal ran back in the opposite direction (Fig. S4). Furthermore, the response was abolished in the dark (Fig. 4, D and E). Thus, widefield cells show clear responses to stimuli displaced across the visual field by the animal’s locomotion. To interrupt the flow of visual information between the SC and LVC we selectively blocked synaptic release in widefield cells by conditionally expressing tetanus toxin light chain (TetTx; Fig. 5A), a perturbation that was previously shown to strongly reduce visual responses in LVC (*18*). After a ten-day recovery period, TetTx-injected mice were trained and tested on the linear track, as in previous cohorts (Fig. 5B).

**Fig. 4.**
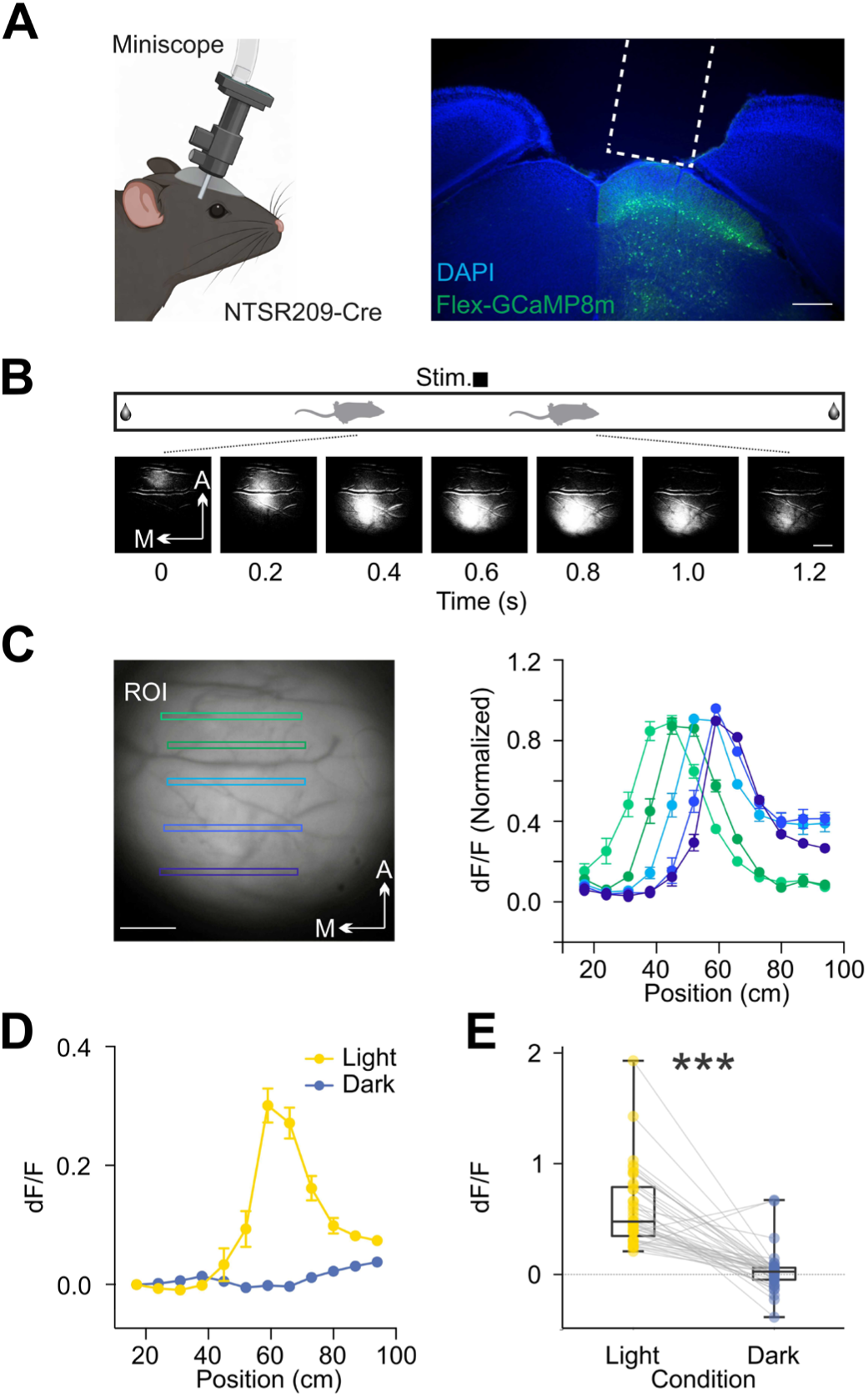
SC Widefield neurons respond to static visual stimuli during locomotion. **(A)** Left: Schematic of the experimental setup showing a Miniscope attached to the head of an NTSR1-GN209-Cre mouse. Right: Representative coronal section showing Cre-dependent expression of GCaMP8m (green) in the superficial layers of the superior colliculus (SC), with DAPI in blue. The white dashed box indicates the approximate position of the GRIN lens. Scale bar: 400μm. **(B)** Top: Schematic of the linear track, with a static visual stimulus (black square) positioned along the side. Bottom: Time-lapse of a calcium activity in neuropil of widefield cells of the SC recorded through the Miniscope during a representative traversal of the track. M: Medial, A: Anterior. Scale bar: 200μm. **(C)** Left: Representative field of view with color-coded Regions of Interest (ROIs) drawn along the anterior-posterior axis of the SC. Right: Normalized fluorescence changes (dF/F) for the corresponding ROIs plotted against the animal’s position on the track (cm). Note the sequential activation of ROIs correlating with spatial position. Scale bar: 200μm. **(D)** Average calcium response (dF/F) relative to track position during Light (yellow) and Dark (blue) conditions. Error bars indicate SEM. n = 10 trials. **(E)** Pairwise comparison of peak dF/F responses in Light versus Dark conditions across mice. n = 40 trials from N = 4 mice; ***, p = 2.6325e-07 (Wilcoxon signed-rank test).

**Fig. 5:**
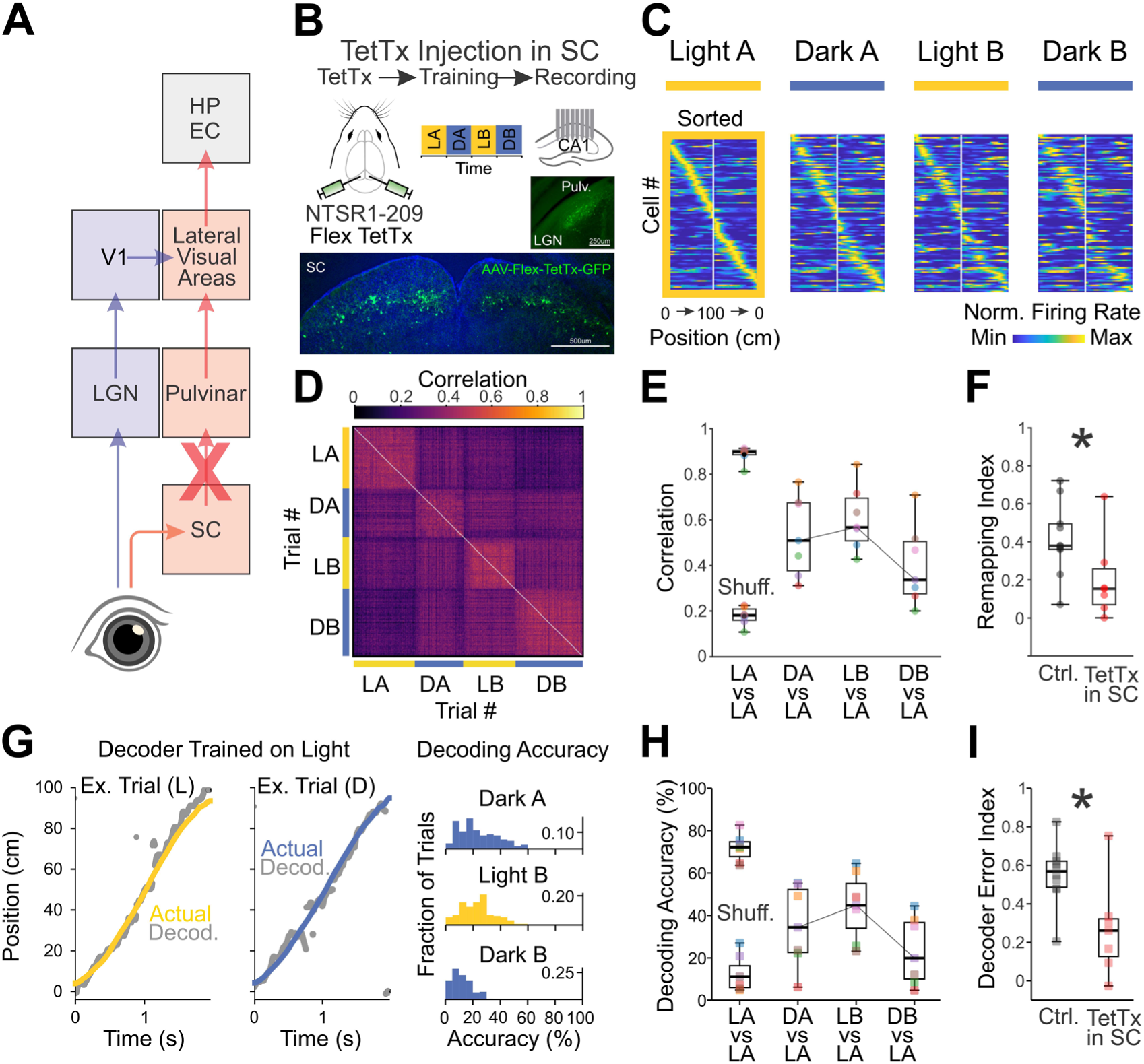
Silencing SC widefield neurons consolidates light and dark spatial maps. **(A)** Schematic of visual pathways, with silencing of the superior colliculus (SC) projection to the pulvinar indicated by the red X. **(B)** Experimental paradigm and histology. Top: Timeline with AAV-Flex-TetTx-GFP injection in SC prior to training and recording. Bottom: Representative micrographs showing viral expression (green) in the SC (injection site) and in SC projections to the pulvinar, but not LGN, following injection of AAV-Flex-TetTx-GFP into an NTSR1-GN209-Cre mouse. **(C)** Normalized firing rate maps of CA1 place cells from a representative SC-pathway silenced mouse, sorted by peak firing position in Light A. Maps represent concatenated leftward and rightward runs. **(D)** Pairwise Spearman correlation matrix of population activity across all trials for a representative SC-pathway silenced mouse (chronologically ordered). **(E)** Spearman correlation of population vectors between odd and even trials within and across blocks for all SC pathway silenced mice (N = 7 mice). “Shuff.” (Shuffle) represents correlations between population vectors in odd trials and spatially randomized population vectors in even trials. Box plots indicate median (center line), interquartile range (box edges), and range (whiskers). **(F)** Remapping Index comparing Control (Ctrl., data from Figure 1) and SC pathway silenced (TetTx in SC) mice. Index is calculated as in Figure 1, with lower values indicating reduced remapping. N = 7 SC pathway silenced mice; N = 10 control mice; *, p = 0.033 (Wilcoxon rank-sum test). **(G)** Left: Bayesian decoding of position in a SC pathway silenced mouse using a decoder trained on Light A. Plots show actual position (colored lines) versus decoded position (grey points) for a light and dark example trial. Right: Distribution of decoding accuracy for Dark A, Light B, and Dark B trials using the Light A decoder. **(H)** Decoding accuracy across all trials for each SC pathway silenced mouse (N = 7 mice). Shuffle was calculated by randomly permuting decoded positions across spatial bins and calculating the percentage of bins in which the error fell within the actual Light A threshold. **(I)** Decoder Error Index comparing Control and SC pathway silenced mice, as calculated in Figure 1. N = 7 SC pathway silenced mice; N = 10 control mice; *, p = 0.025 (Wilcoxon rank-sum test).

TetTx-injected mice formed place fields that tiled the track, exhibited similar average SMIs in both light and dark (Fig. S2D), and allowed for accurate decoding of the animal’s position. Strikingly, however, changing from light to dark had only a minor effect on the hippocampal representation of the track, thus recapitulating the effect of LVC lesions. In TetTx-injected mice the remapping index was significantly lower than in control animals (Fig. 5, C to F; Remapping Index, TetTx: Median = 0.151, IQR = 0.1888, N = 7, p = 0.033, Control vs. TetTx), and in contrast to control animals, the SMI of place cells did not change from light to dark (Light Median SMI = 6.40, IQR = 2.2646; Dark Median SMI = 6.45, IQR = 3.5768; N = 7; p = 0.938). Crucially, in TetTx-injected animals, a decoder, trained on light trials, was significantly more accurate at decoding the animal’s position in darkness as compared to control mice (Fig. 5, G to I; Decoder Error Index TetTx: Median = 0.261, IQR = 0.188, N = 7, p = 0.031, Control vs. TetTx). These data indicate that visual information conveyed by the SC via widefield cells can shape spatial maps of the track in the hippocampus.

## Discussion

This study demonstrates that LVC is an essential node in the transmission of visual signals to the hippocampus and that these signals are conveyed to LVC via the SC through a specific neuronal subtype, the widefield cell. LVC sends axonal projections to the entorhinal cortex, a structure upstream of the hippocampus, and to the hippocampus itself (*13*, *24–26*). As such, it is well-suited to provide visual signals to place cells. Indeed, prior anatomical studies have suggested that LVC may represent a gateway of visual information to the hippocampus, though other routes from medial visual cortex have also been proposed (*13*, *25*).

V1 has been generally considered the main relay of visual information to the hippocampus, largely because it was presumed to be the primary entry point for visual signals to cortex (*9*). However, visually-evoked responses in LVC rely principally on the SC rather than V1, positioning the LVC as an alternative, V1-independent gateway for visual information to reach the cortex and, subsequently, the hippocampus (*16*, *18*). Consistent with this proposed function, light-dark remapping–which depends on LVC–was disrupted by SC widefield cell silencing but left intact following V1 lesions. While this study does not exclude the involvement of V1 under different circumstances, such as when spatial maps may depend on the fine discrimination of visual landmarks, it establishes the SC pathway as a main route of visual signals for hippocampal representations of space.

Widefield cells are the main conduit for visual information from the SC to LVC, information which is relayed via the pulvinar (*16–18*). These neurons, which receive direct input from the retina, are uniquely sensitive to both the motion of stimuli across the visual field and to the visual expansion of objects (*19*, *27*). Although these dynamic visual properties have usually been studied in the context of motion of external objects (*28*, *29*), they can also be generated by the movement of the animal itself relative to static objects in the environment, as shown here (Fig. 4). Visual flow, as generated by forward locomotion, indeed supports path integration in the hippocampal representation of space (*6*, *30–33*). We thus hypothesize that by signaling the animal’s movement relative to stationary external objects, widefield cells contribute to the representation of the animal’s position in the hippocampus. Other features of the visual scene may also be captured by widefield cells and contribute to spatial representation in the hippocampus (*34*).

The observed remapping between light and dark was more substantial than previously reported (*8, 35, 36*). We ensured very low light levels (<100 R*/rod/s), which may not have been achieved in previous studies in the absence of special precautions. Furthermore, we familiarized the animals in alternating blocks of both light and dark over several days prior to our recordings. This familiarization protocol may have further contributed to the formation of independent maps under two distinct lighting conditions, analogous to the reported formation of directionally selective place fields following the repeated traversal of the same trajectory in opposite directions (*8*, *37*). Finally, we also ensured maximal stability of the experimental context between the familiarization and the recording periods. For this reason, all lesions were performed at least ten days before the beginning of the familiarization period. Furthermore, we avoided any acute perturbation, optogenetic, chemogenetic or surgical, between the familiarization and recording period, or during the recording period, which, by changing the experimental context, can per se lead to profound remapping (*38*).

Earlier work has placed the hippocampus at the top of a visual processing hierarchy, starting from the retina and passing through V1 (*9*). This study demonstrates that another important branch of the vertebrate’s visual system, namely the SC pathway, conveys visual information to the hippocampus, independently of V1. The SC contributes to blindsight, the ability of patients who are cortically blind, for example as a consequence of V1 lesions, to use visual information to guide behavior in the absence of visual awareness (*39–41*). Some of these patients retain the ability to navigate (*42*). The SC pathway to the hippocampus described here may mediate this residual capacity.

## Acknowledgments

We thank Pooja Saraf, Qui Ying Wu and Leo Ruan for technical assistance, Loren Frank and Eric Denovellis for sharing their decoding method and the former and current members of the Scanziani lab for discussions. We thank Roger A. Nicoll and Loren Frank for comments on the manuscript. Gemini 3.1 Pro was used to draft portions of the materials and methods and figure legends, with the authors’ notes and code used as input. All such sections were fully reviewed and edited by the authors.

## Funding

National Institutes of Health grant R01EY025668 (MS)

Howard Hughes Medical Institute (MS)

National Institutes of Health grant NEI 1F31EY033661-01 (JMB)

National Institutes of Health grant K99EY033850 (YS)

National Institutes of Health grant R00EY033850 (YS)

Japan Society for the Promotion of Science (YS)

Wellcome Trust CDA 225353/Z/22/Z (SR)

Wellcome Trust and the Royal Society 222583/Z/21/Z (RB)

Helen Hay Whitney Fellowship (JMR)

NOMIS Foundation Fellowship Program at the Gladstone Institutes (YV)

## Author contributions

JMB, RB and MS designed the study.

JMB, RB, and SR conducted all electrophysiological experiments and performed the analysis.

CW performed decoding analysis and profilometry.

J.R. conducted miniscope imaging and analysis.

YS and CW developed the linear track protocol with chronic hippocampal recordings under light and dark conditions.

YV developed and shared the protocol for the profilometry in the Paz lab.

JMB, RB, SR and MS wrote the manuscript.

All experiments were performed in the Scanziani lab at UCSF.

## Data, code, and materials availability

All data and analyses necessary to understand and assess the conclusions of the manuscript are presented in the main text and in the supplementary materials. Raw data will be made available upon reasonable request.

## Supplementary Materials

### Materials and Methods

#### Animals

All experiments were performed in accordance with protocols approved by the Institutional Animal Care and Use Committee of the University of California, San Francisco. Experimental mice were all male from the C57/B6 background; transgenic lines include NTSR1–GN209–Cre (RRID: MMRRC_030780-UCD). Mice were at least two months old at the time of the experiments. Mice were housed on a reverse light cycle (12/12 hours) with experiments carried out exclusively during the dark cycle.

#### Linear Track (Electrophysiology)

A custom linear track 100 cm in length and 5 cm in width was constructed from a sheet of chemical-resistant ⅛ inch thick styrene plastic (TAP Plastics, San Francisco, CA, USA). To provide somatosensory location cues, 7 distinctly textured equally-sized tiles (14 cm each) were cut from sheets of plastic and sandpaper and glued to the floor of the track. The track was mounted on aluminum posts 50 cm above the surface of a table to prevent the mouse from escaping. Plastic lick-ports were attached at either end of the track, and a custom closed-loop system (Bonsai) was used to track the animal in real time and deliver a soymilk reward whenever the animal completed a run from one end to the other. The entire apparatus was enclosed in a light and soundproof box. To further isolate the mouse from external sound and light cues, the box was surrounded by light-proof curtains and white noise machines were placed at either end of the track. To illuminate the inside of the arena, the track was surrounded by regularly spaced white LED strips controlled by custom code (Matlab) and a microcontroller (Arduino Uno). To allow the mouse to be tracked even in complete darkness, far-infrared LED panels (940 nm LED board, SCS Enterprises) were mounted next to a CMOS camera (Basler acA1300- 200um) above the track. Before recording, mice were food-deprived to 80% of body weight and habituated on the linear track for at least three days, or until they could complete 200 laps in 2 hours, using the same alternating light/dark block protocol as on the day of recording. Pose estimation and tracking were performed post hoc with DeepLabCut (*43*).

Electrophysiological recordings were acquired using an RHD2000 data acquisition system (Intan Technologies). The signal was sampled at 20 kHz and band-pass filtered between 0.1 Hz and 7.5 kHz. To minimize weight and torque on the animal’s head, the headstage was tethered to the recording system using an ultra-thin, flexible cable (Intan Technologies). To maintain unimpeded movement, the tether was routed through a custom 3D-printed active commutator (*44*).

#### Linear Track (Miniscope Imaging)

In a subset of experiments involving superior colliculus miniscope calcium imaging, the linear track was identical to that described above except that the walls were painted with a shade of gray matching the E-Ink screen. Mice were food-deprived to 80% of their free-feeding body weight and habituated to the linear track for at least 3 days; they were then habituated to wearing a dummy miniscope for 2 additional days while performing runs on the track to acclimate to the added weight. On the day of the experiment, mice were fitted with the Miniscope (*23*; UCLA Miniscope; miniscope.org) and its coaxial cable was attached to an Open Ephys Miniscope commutator. The commutator position was controlled via Bonsai commands to prevent cable tangling. Once the optimal field of view was confirmed, recording commenced in Bonsai. A 0.5 in × 0.5 in black square was presented on an E-Ink screen positioned 15 cm from the middle section of the linear track at an elevation of 20° above the animal. Track illumination was dimmed progressively every 10 trials until completely off (“Dark” condition).

#### Light Level Measurements

To estimate the light reaching the retina during dark conditions, we made a series of measurements along the track and used these to calculate photoisomerizations per rod per second. First, the spectral distribution of the infrared (IR) light sources (940 nm LED board, SCS Enterprises) was measured using an SM240 Spectrometer and SM32Pro software (SpectralProducts). Total light power during dark conditions was then measured using a Model S471 Optometer equipped with a Model 268R Flat Response Radiometric Sensor (1 cm² active area; UDT Instruments). Power measurements were taken at 18 locations within the arena: three track positions (left, center, and right) crossed with six sensor orientations at each position (ceiling wall, 45° wall, lateral wall, ceiling curtain, 45° curtain, and lateral curtain). Using the spectral distribution and power readings, rod photoisomerizations at each orientation were calculated using the CalibrateLight software package (*45*). The final reported value (<100 R*/rod/s) was calculated from the average across the lateral, 45°, and ceiling orientations.

#### Surgeries

Mice were anesthetized in an induction chamber under 4% isoflurane in oxygen. Following induction, mice were secured in a stereotaxic frame (Kopf) and maintained under 2% isoflurane in oxygen to maintain anesthesia for the duration of the surgery. A protective petrolatum ophthalmic ointment (Puralube) was applied to the eyes. The surgical area was sterilized with saline, alcohol, and povidone-iodine (Betadine). For analgesia, a 2% lidocaine cream (Akorn Pharmaceutical) was topically applied to the incision site, while buprenorphine (0.1 mg/kg) and carprofen (5 mg/kg) were delivered via subcutaneous injection. These subcutaneous analgesics were re-administered at 6, 12, and 24 hours postoperatively.

##### Probe Implantation

Mice underwent a two-stage surgical procedure consisting of headplate attachment followed by silicon probe implantation. The hardware was constructed based on a previously described recoverable microdrive system (*46*). During the first stage, the baseplate was securely affixed to the skull using a combination of dental adhesive (OptiBond Universal, Kerr Dental) and dental cement (Unifast LC, GC America). For the probe implantation, the surgical site was stereotaxically marked, and a 1.2 mm long craniotomy was made at a 45-degree angle to the midline, centered at anteroposterior (AP) −2.0 mm and mediolateral (ML) 1.5 mm from bregma. A 128-channel silicon probe (128-8, Diagnostic Biochips) was subsequently lowered into the brain to an initial depth of 0.8 mm from the cortical surface. The exposed brain tissue within the craniotomy was then sealed with a silicone elastomer (Kwik-Sil, World Precision Instruments), and the microdrive holder was permanently secured to the headplate using C&B Metabond and a light-curing dental adhesive (Triad). The day prior to the first recording session, the probe was advanced to its final target depth of approximately 1.2 mm. Proper placement within the CA1 pyramidal layer was confirmed online by identifying characteristic electrophysiological signatures and histologically post hoc.

##### VC lesions

Focal cortical lesions were targeted to the primary visual cortex (V1) or to lateral higher visual areas (HVAs) using stereotaxic coordinates derived from the Allen Mouse Brain Common Coordinate Framework (*22*) (CCFv3). Mice were prepared for surgery as described above. The skull surface was cleaned and freed of connective tissue to reveal cranial landmarks (bregma and lambda). Using these landmarks as reference, the boundaries of the lesion area were drawn on the skull according to CCF-derived coordinates. Relative to bregma (in mm), the specific AP coordinates and their corresponding mediolateral (ML) ranges were as follows: AP −2.47, ML 2.20 to 2.80; AP −2.77, ML 2.01 to 3.15; AP −3.07, ML 1.80 to 3.40; AP −3.37, ML 1.68 to 3.46; AP −3.67, ML 1.57 to 3.48; AP −3.97, ML 1.39 to 3.50; and AP −4.27, ML 1.39 to 3.43.

For lateral HVAs, the coordinates were centered on POR, extending from –3.75 mm to –4.87 mm posterior to bregma and 3.7–4.8 mm lateral. A craniotomy was made with a Gesswein microburr attached to a Foredom dental drill by following the drawn outline, and cortical tissue within the delineated area was surgically removed by microdissection with a fine microknife and gentle suction through a pipette. The exposed cortical surface was sealed with a thin layer of Kwik-Cast (World Precision Instruments) and covered with dental cement to ensure long-term protection of the lesion site. Animals were allowed to recover for at least ten days before behavioral experiments. Lesion extent and specificity were verified post hoc by histological reconstruction of coronal sections aligned to the Allen CCF and profilometry, confirming localized ablation of V1 or the lateral HVAs (LI, POR).

##### Tetanus Toxin Light Chain Viral Injections

Six craniotomies, each measuring 50 μm, were drilled using a Gesswein microburr attached to a Foredom dental drill. The viral preparation (AAV.2.1.flex.TeLC.GFP diluted to 5 × 10^12^ copies/ml in PBS) was drawn into a glass pipette that had been mechanically pulled and beveled to a tip diameter of 20μm. At each of the six injection locations (AP −3.20 mm, ML ±0.70 mm, DV −1.30 mm rising to −1.50 mm over the course of the injection; AP −3.80 mm, ML ±0.95 mm, DV −1.20 mm rising to −1.10 mm over the course of the injection; AP −4.40 mm, ML ±0.70 mm, DV −0.70 mm rising to −0.80 mm over the course of the injection), 400 nL of virus solution was delivered via a micropump (UMP-3, WPI) at a rate of 80 nl per minute. The pipette remained in position at the delivery site for 20 minutes following the injection, after which it was gradually withdrawn and the surgical incision closed with sutures. Experiments were performed at least ten days after the injection to ensure adequate viral expression. After the experiment, mice were perfused, and their brains were sectioned to confirm the injection site and viral expression histologically.

##### GCaMP Viral Injection and GRIN Lens Implantation

For superior colliculus calcium imaging experiments, mice underwent two serial surgical procedures spaced 4 weeks apart. During the first surgery, mice were prepared as described above. A 1 mm diameter craniotomy was made above the superior colliculus on the right hemisphere (centered at AP −3.6 mm, ML +0.7 mm from bregma). Then, 400 nL of AAV1-hSyn-FLEX-Soma-GCaMP8m (diluted to ∼1 × 10¹³ genome copies/ml) was injected into the superior colliculus on the right hemisphere (AP −3.6 mm, ML +0.7 mm, DV −1.7 mm) of NTSR1–GN209–Cre mice. After the pipette was removed, the mouse remained on the stereotaxic frame for 20 min to allow complete diffusion of the virus. The cortex below the craniotomy (consisting of the medial higher visual areas) was aspirated with a 27-gauge blunt syringe needle attached to a vacuum pump while constantly being irrigated with ACSF. Once the superior colliculus became visible, bleeding was controlled using surgical foam (Surgifoam). A 1mm diameter × 4mm length GRIN lens (CLHS100GFT003, GoFoton) was then slowly lowered into the craniotomy using a stereotaxic holder (XCL, Thorlabs). The lens was fixed with Norland Optical Adhesive, and dental acrylic was applied to cement the implant in place and to cover the remaining exposed skull. The top of the exposed lens was covered with Kwik-Sil (World Precision Instruments) to protect it, and the Kwik-Sil was covered with dental cement.

##### Baseplate Implantation

Four weeks after GRIN lens implantation, mice were re-anesthetized to attach the miniscope baseplate. The overlying dental cement was drilled off and the Kwik-Sil was removed to reveal the top of the lens. A 3D-printed headplate for head-fixation experiments was cemented in place. The miniscope (with attached baseplate) was lowered near the implanted lens while the field of view was monitored in real time. The miniscope was rotated until a well-exposed field of view was obtained, at which point the baseplate was fixed to the implant with dental cement.

#### Profilometry and Histology

For the anatomical analyses described below, brain tissue was prepared by transcardially perfusing mice with 10 mL of phosphate-buffered saline (PBS; chilled to 4°C), followed by 10 mL of 4% paraformaldehyde (PFA) in PBS. Brains were then extracted and submerged in a 4% PFA solution for overnight fixation at 4°C.

##### Profilometry Analysis

Prior to sectioning, whole-brain surface profiles were obtained from the dorsal surface using a 3D laser-scanning confocal microscope (Keyence VK-3000, QB3 Center UC Berkeley). Brains with lateral visual area lesions were imaged using a 3D-printed 30° wedge to expose the posterior lateral lesion. Lesion volume and surface area of the missing cortex (i.e. cross-sectional area) in each hemisphere were calculated using MultiFileAnalyzer software (Keyence), where volumes were defined as the space between a reference cortical plane and the lesion profile. To approximate the missing cortical surface, the reference plane was determined by selecting a series of points on the cortex adjacent to the lesion. Final volumes and surface areas are presented as averages across hemispheres (Supp. Fig. 3). 3 LVC lesioned and 4 V1 lesioned brains were processed for histology before the profilometry protocol was established, and thus were not included with this analysis.

##### Lesion Quantification by Histology

After optical 3D profilometry (above), brains from lesioned mice were sectioned to 100 μm thickness with a vibratome and mounted on slides with Vectashield mounting medium with DAPI (Vector Laboratories H1500). Sequential micrographs were acquired with an Olympus MVX10 MacroView microscope. Micrographs in which the lesion was present were loaded into FIJI (ImageJ) and aligned to the Allen Common Coordinate Framework. Lesion coordinates were then manually measured, and the lesion area was quantified for each micrograph. Lesion volume was estimated by interpolating the lesion area between sections.

##### Lesion Mapping and Visualization

Brain region boundaries for visual cortical areas (AM, A, PM, V1, RL, L, AL, LI, POR) were defined using the Allen Common Coordinate Framework. The extent and position of each lesion in the medio-lateral and antero-posterior axes were measured in each mouse as described above. ‘Median’ lesion extent was determined by calculating the median position of the lateral and medial edges at each bregma section covered by the lesions and interpolating between sections. Median lesions were then overlaid on the individual lesions and the Allen-defined visual cortical area boundaries.

#### Electrophysiology Analysis

##### Preprocessing

Electrophysiological data were sorted using Kilosort 2 (47) and manually curated with Phy (48). A filter of 1 refractory period violation per 500 spikes was used to remove contaminated units. Videos taken during recordings were processed with DeepLabCut to track the mouse’s position along the track throughout the course of the recording; position stamps for ‘left’ and ‘right’ ears were averaged to calculate the mouse’s position in each frame. Timestamps for each frame were read out as TTL pulses from the camera and sent to the Intan recording system (Intan RHD2000); these timestamps were used to assign frames spikes from each unit to the corresponding video frames. An additional TTL was sent to the Intan system from the camera to indicate the onset and offset of the video recording, with a third TTL from the MATLAB system controlling the lights via a National Instruments Data Acquisition Board (NI USB-6001) to indicate the lighting condition of the track, and thus assign both frames of mouse behavior and unit spikes to ‘light’ or ‘dark’ epochs. Additionally, a TTL through a photodiode was used to continuously readout light levels and ensure that actual light conditions in the track corresponded to intended light conditions.

To reconstruct neural maps of the track, we identified epochs during which the mouse was running from one end of the track to the other. These were defined as periods in which the mouse left one lickport and approached the other lickport no more than 10 seconds later. The track was split into spatial bins of 5 cm and a minimum run period of 1 second was used to filter artifactual ‘jumps’ in the tracking data. Each valid run was then assigned to a ‘light’ or ‘dark’ condition based on the lighting TTL status from MATLAB. Any runs that spanned a change in lighting were discarded. For each run, spike timestamps from each unit were loaded from Phy and assigned to the corresponding spatial bin based on the mouse’s position at that timestamp.

##### Place Cell Analysis

Spatial firing fields were identified from position-binned firing rate maps computed across all traversals of the linear track. The track was initially divided into 20 spatial bins, each 5 cm in length. For each neuron, the spatial firing rate was calculated as the number of spikes recorded within each bin divided by the total occupancy time in that bin. The first and last spatial bins were excluded from analysis because these regions corresponded to the locations of two reward ports, where animals frequently paused or lingered to collect the reward, potentially introducing biases in firing rate estimates. After this exclusion, data from both running directions of the track were merged, yielding a combined linear track of 36 bins. Peaks in the resulting rate map were detected using MATLAB’s *findpeaks* function with a minimum peak firing rate of 1 Hz and a minimum peak width of 5 cm. For each detected peak, the boundaries of the spatial firing field were defined as the contiguous bins where firing remained above 40% of the peak rate. Field edges were determined by extending left and right from the peak until the rate fell below this threshold. Fields narrower than 5 cm or broader than 50 cm were excluded. Neurons were classified as place cells if they contained at least one spatial field meeting these criteria. For each field, we extracted the start and end positions, width, peak firing rate, and peak location.

Place field statistics (width, firing rate, Spatial Modulation Index (SMI)) were computed from grouped population firing rate matrices for control, laterally lesioned, V1 lesioned, and SC pathway silenced mice, organized by light and dark conditions. To calculate width, firing rates were normalized for each unit and condition, and field widths were defined as the number of contiguous spatial bins that exceeded 40% of the peak firing rate. Since some units had place fields in one condition but not the other, only conditions with peaks at least two standard deviations above baseline were included. Peak firing rates were calculated as the maximum firing rate across spatial bins for each unit in light or dark.

SMI was calculated for each unit and running direction as a normalized version of the spatial information content (*8*) (SIc). To normalize the SIc, we circularly shuffled a cell’s spiking activity across the track 100 times and computed the SIc for each shuffle. From this distribution, we obtained the mean (m) and standard deviation (s). SMI was defined as the z-score of the actual SIc value: SMI = (SIc actual - m)/s (*49*).

##### Epoch Selection

Prior to computation of place fields and population analyses, blocks of luminance conditions (Light A, Light B, Dark A, and Dark B) were screened for inclusion using pre-established criteria. Because our primary analysis tested the null hypothesis that light-dark remapping remains unchanged following a given perturbation, we designed these criteria to conservatively select the most stable epochs for each mouse. The inclusion criteria applied to the population of identified place cells were: (1) the Spearman auto-correlation between firing-rate maps derived from odd versus even trials within each individual block exceeded 0.85; (2) the correlation between blocks A and B within the same luminance condition exceeded 0.3; and (3) the session contained a minimum of 35 place cells. When multiple block sets satisfied all three criteria, the set with the highest average intra-luminance correlation (the mean of the Light A–Light B and the Dark A–Dark B correlations) was selected. These stability criteria were applied uniformly across all experimental groups. Light-dark remapping was still evident across groups even without the final epoch selection step.

##### Remapping Index

To quantify spatial stability, population vector correlations were calculated by comparing matrices of place cell firing rates from odd trials of a reference condition against even trials of all other conditions. Using the average population vector correlation for each pair of conditions, a remapping index for each mouse was calculated as the difference between the same-condition Light A–Light B correlation and the average cross-condition correlation (Light A–Dark A and Light A–Dark B). This value was normalized by the dynamic range between the internal self-correlation of Light A (Light A even vs. Light A odd trials) and chance (Light A shuffle):

*RI = (Corr_LA,LB_ − (Corr_LA,DA_ + Corr_LA,DB_) / 2) / (Corr_LA even, LA odd_ − Corr_LA shuffle_)*

Light A shuffle was calculated by randomly permuting the firing rates across spatial bins 1,000 times per trial in the Light A condition, calculating the Spearman correlation coefficient, and averaging across shuffles and trials to obtain a null distribution.

##### Population Correlation

“Checkerboard” correlation heatmaps were generated by concatenating firing rate matrices of place cells from leftward and rightward linear track traversals on a trial-by-trial basis. The 2D Spearman correlation coefficient was calculated between these firing rate matrices in chronological order for every possible pair of trials. The resulting matrix was visualized as a heatmap with Matplotlib using the ‘inferno’ colormap, normalized between 0 and 1.

##### Decoder Analysis

Spatial position from hippocampal spiking activity was decoded using the Bayesian state space algorithm from Denovellis et al. 2021 (*20*). Briefly, for a given luminance condition (e.g., Light A), an encoding model was built to relate the spiking from all sorted CA1 cells in the recording to the animal’s position in 1ms bins using trial period data from the first 80% of time in that condition. We then decoded spatial position from CA1 spiking activity in 0.5cm position bins and 1ms temporal bins across each ‘test’ luminance condition trial (i.e., ‘held out’ trials in the remaining 20% of time in Light A and all trials in Dark A, Light B, and Dark B). The state space model had two states, continuous and fragmented, modeled using a random-walk or uniform transition matrix, respectively. The model output was the posterior probability of position across the two states. The ‘decoded position’ in each time bin was determined as the position of the peak of the posterior probability distribution in the continuous state. Importantly, separate models were trained for each running direction on the linear track and the decoding results were concatenated and sorted in time for analysis.

##### Decoding Accuracy

We quantified place field reorganization across luminance conditions by calculating a ‘decoding accuracy’ metric, defined as the average proportion of position bins across trials whose decoding error (the absolute difference between the animal’s decoded and actual positions) fell within the average Light A decoding error (i.e., the average decoding error across the ‘held out’ trials in Light A). Since the encoding model was trained on data from Light A, the average Light A decoding errors were typically low (within 5–10 cm). Accordingly, luminance conditions in which the decoded position closely tracked the animal’s actual position (i.e., conditions with similar place field maps compared to the Light A condition) yielded high decoding accuracy. Conversely, luminance conditions in which the decoded position was widely deviated from the animal’s actual position (i.e., conditions with dissimilar place field maps compared to Light A condition) yielded low decoding accuracy.

##### Decoder Error Index

Using the average decoding accuracy value for each luminance test condition, a decoder error index for each mouse was calculated as the difference between the accuracy of the decoder in Light B and the average of its accuracy in Dark A and Dark B, normalized by baseline range in the accuracy between Light A and the Light A shuffle:

*RI = (Light B − (Dark A + Dark B) / 2) / (Light A − Light A shuffle)*

The Light A shuffle was calculated by randomly permuting the decoded positions across spatial bins 1,000 times per trial in the Light A test condition, calculating the percentage of bins whose decoding error fell within the actual Light A error threshold, and averaging across shuffles and trials to obtain a null distribution.

#### Miniscope Video Analysis

Miniscope data were preprocessed by manually selecting a region of interest (ROI) in the video corresponding to calcium fluctuations driven by the visual stimulus as it moved through the nasal-temporal axis of the mouse’s visual field. Raw pixel intensity within this ROI was averaged at each time point and subtracted by the average intensity from a sub-ROI placed at the far rightmost position of the GRIN lens (to correct for global fluctuations in the imaging window unrelated to the stimulus). Data were aligned to behavior using custom MATLAB scripts that synchronized calcium signals with the animal’s position relative to the visual stimulus. ΔF/F was calculated for each complete traversal of the linear track from one end to the other. The track was divided into 12 equally spaced spatial bins, and the ΔF/F signal was averaged within each bin.

#### Behavioral Analysis

To compare locomotor behavior between light and dark conditions, the mouse’s position across every frame was tracked with DeepLabCut and periods corresponding to leftward and rightward runs were identified as described above. To reconstruct periods containing complete laps of the track, sequential leftward and rightward runs were concatenated. Unmatched runs or laps that occurred partly in light and partly in darkness were discarded. The trajectory of a lap was then defined by splitting the track into 100 bins along its length and calculating the mouse’s median position within each bin’s width. The median trajectory was then calculated from the trajectories of all laps for each condition. Trajectories were segmented into 30 bins for display and visualized with the ‘viridis’ colormap for velocities and ‘magma’ for accelerations (matplotlib). Instantaneous velocity was calculated as the Euclidean distance between mouse center-points in sequential frames of video, scaled by frames/second. Average velocity was then calculated for each lap and for all laps in each condition. Instantaneous accelerations were derived from differences in downsampled velocities to account for jitter. Average acceleration was then calculated for each lap and for all laps in each condition.

To isolate the effects of the track’s lighting condition from trial-to-trial variations in locomotor activity of the mouse, we matched individual trials from light and dark epochs based on kinematic similarity. For every possible light-dark trial pairing, a composite “similarity score” was calculated. This score integrated two normalized metrics: the mean absolute spatial deviation between the two trajectories and the absolute difference in the median running velocity across each trial. A subset of trial pairs scoring the highest in similarity was selected for subsequent comparative analysis (Fig. S1), ensuring that differences in neural maps between lighting conditions were not confounded by differences in running speed or path selection.

#### AI Statement

Gemini 3.1 Pro was used to assist in writing and editing sections of code used for analysis. All such code was fully reviewed and validated by the authors.

---

**Fig. S1.**
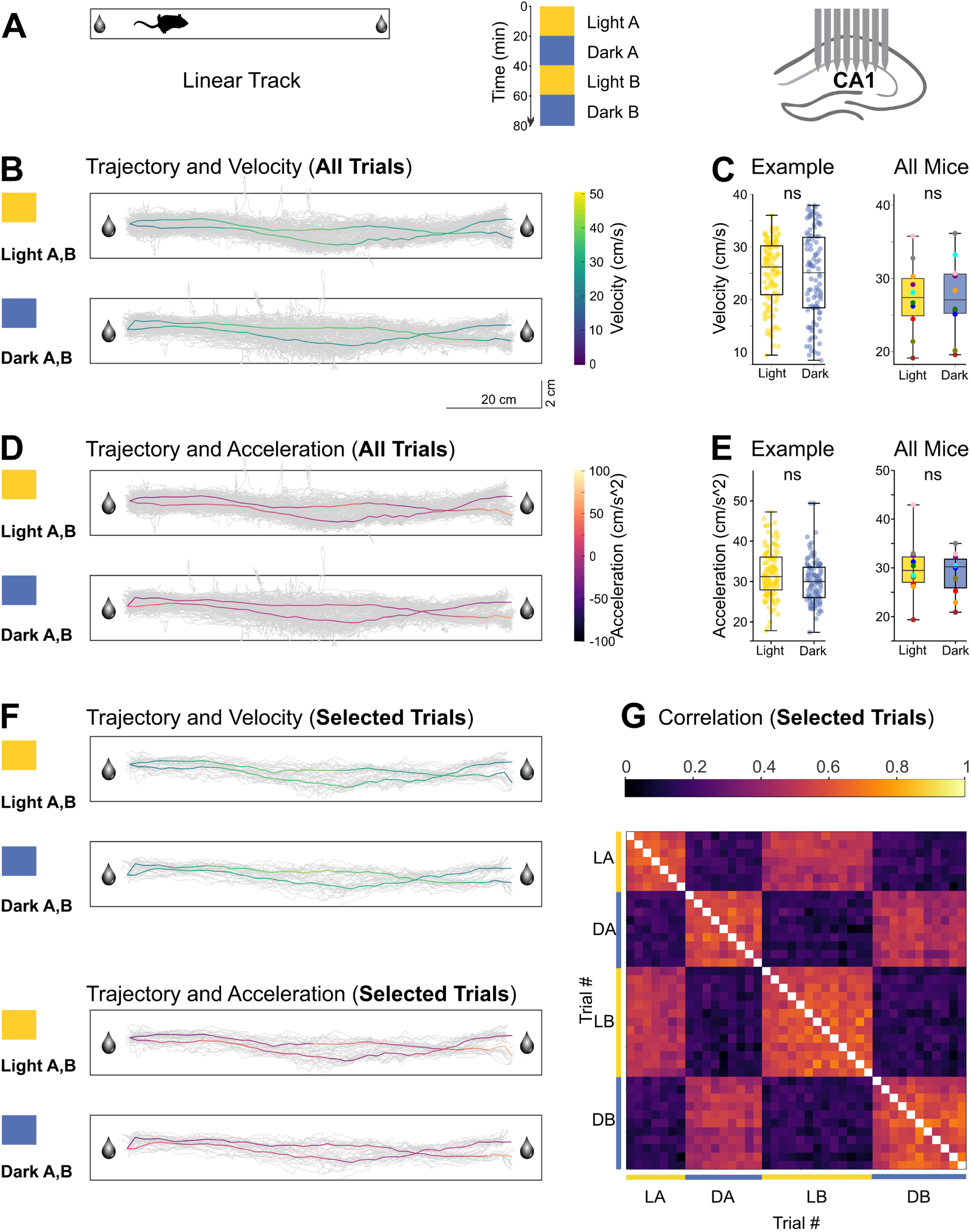
Light-Dark remapping does not depend on behavioral variability. **(A)** Schematic of the linear track task (left), experimental timeline showing interleaved light and dark epochs (center), and the recording site in hippocampal CA1 (right). **(B)** Spatial trajectories for Light and Dark conditions. Individual runs are shown in gray. The median trajectory is overlaid and color-coded by velocity. **(C)** Median velocity in Light (yellow) and Dark (blue) epochs for a representative mouse (left; dots represent individual trials) and the population (right; dots represent individual mice). Box plots indicate median and interquartile range. *ns*, not significant (Wilcoxon rank-sum test for single mouse; Wilcoxon signed-rank test for population). **(D)** Spatial trajectories formatted as in (B), but color-coded by acceleration. **(E)** Mean acceleration in Light and Dark epochs, formatted as in (C). **(F)** Trajectories of trials selected for kinematic similarity (see Methods), color-coded by velocity (top pair) and acceleration (bottom pair). **(G)** Pairwise Spearman correlation matrix of spatial population activity from the kinematically matched trials. Trials are arranged chronologically. Labels indicate epochs: LA (Light A), DA (Dark A), LB (Light B), and DB (Dark B).

**Fig. S2.**
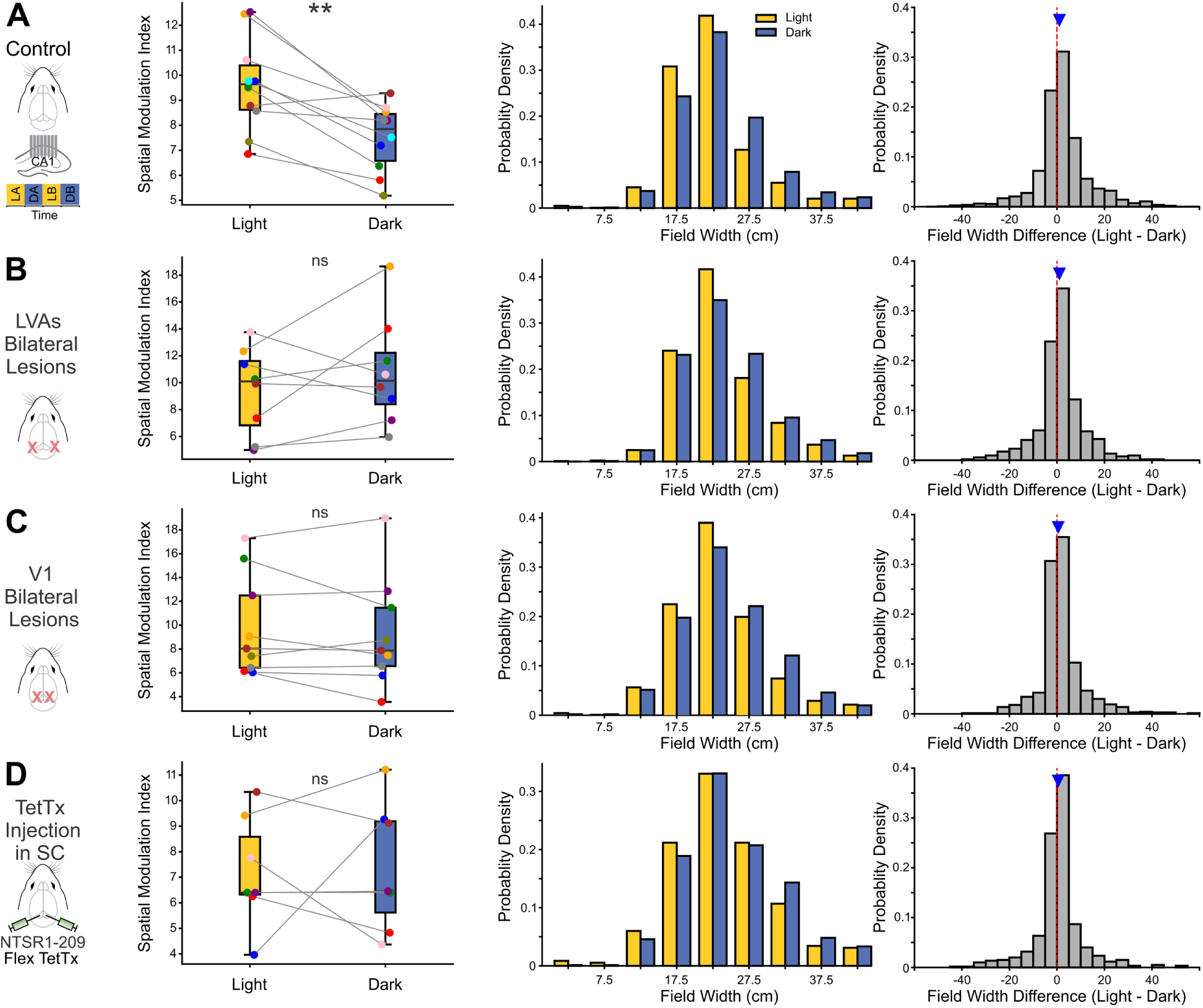
Stability of place field properties across visual pathway manipulations. **(A)** Control mice. Left: Experimental paradigm and median Spatial Modulation Index (SMI) in Light and Dark epochs. Paired lines connect data from individual mice; box plots indicate median and interquartile range. p = 0.006 (Wilcoxon signed-rank test), N = 10 mice. Center: Distribution of place field widths (pooled across mice) in Light (yellow) and Dark (blue) conditions. Right: Distribution of the difference in place cell firing rate (Light − Dark). The blue triangle indicates the population median; the vertical line at 0 indicates no difference. **(B)** Same as (A), for mice with lateral visual cortex lesions. ns, not significant; p = 0.461 (Wilcoxon signed-rank test), N = 8 mice. **(C)** Same as (A), for mice with primary visual cortex (V1) lesions. p = 0.570 (Wilcoxon signed-rank test), N = 9 mice. **(D)** Same as (A), for mice with SC pathway silencing. p = 0.938 (Wilcoxon signed-rank test), N = 7 mice.

**Fig. S3.**
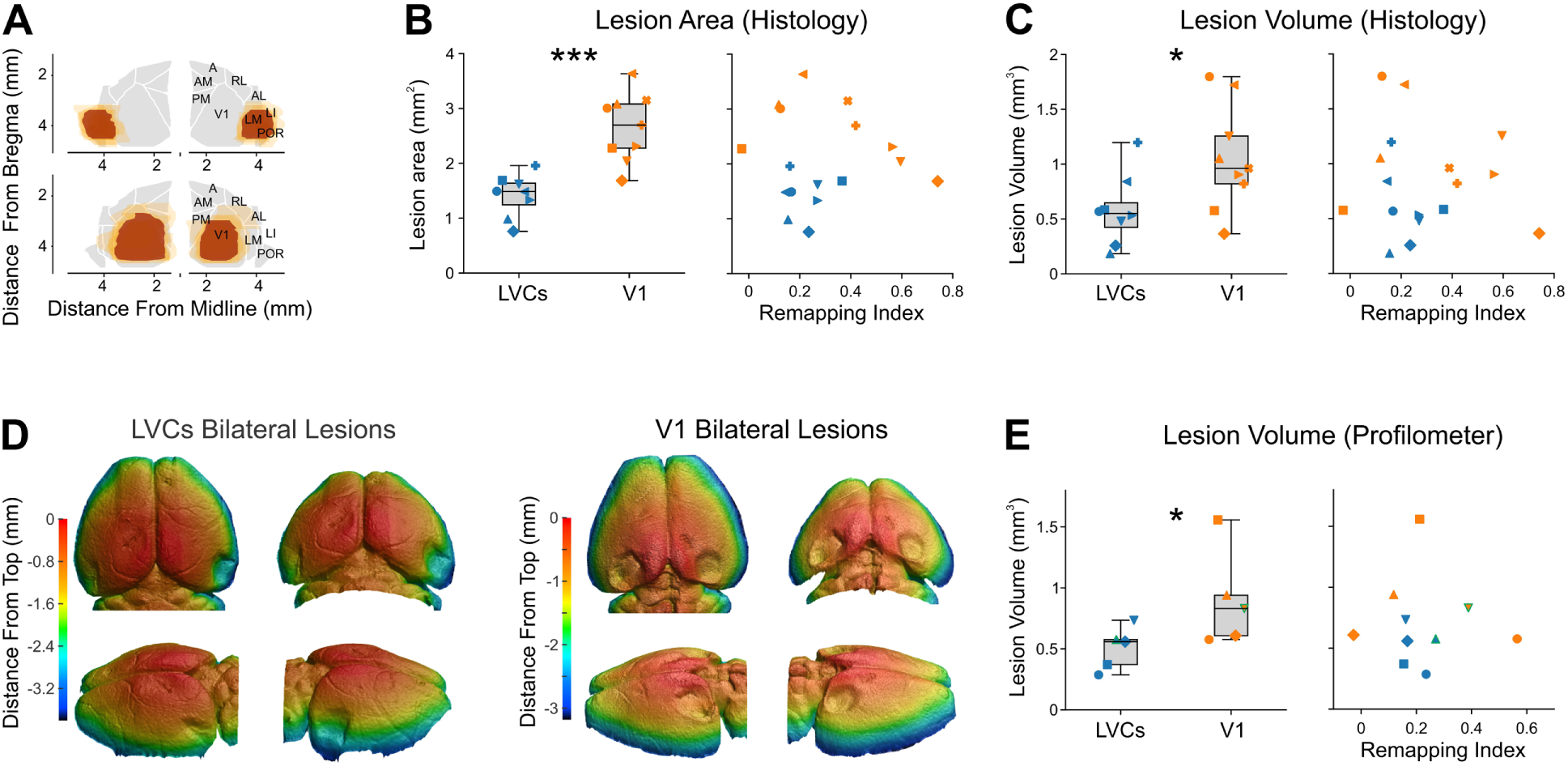
Histological and laser-profilometry analysis of LVCs and V1 lesions. **(A)** Reconstructions from histological data of bilateral LVCs (top, 8 mice) and V1 lesions (bottom, 9 mice) superimposed on a reference map of the visual cortex generated from the Allen Common Coordinate Framework. Dark red indicates the median lesion extent across all mice, overlaid on individual lesion extents (lighter orange). **(B)** Left: Comparison of the areas of LVC and V1 lesions, quantified from histological sections. Box plots indicate median (center line), interquartile range (box edges), and range (whiskers). N = 8 LVC mice, 9 V1 mice; ***, p = 3.29 e-04 (Wilcoxon rank-sum test). Right: Scatter plot correlating V1 and LVC lesion areas with Remapping Index. R = −0.023; p = 0.929 (Pearson’s correlation test); ns, not significant. **(C)** Same as in (B) but for the volumes of LVC and V1 lesions measured from histological sections. Left: *, p = 0.036 (Wilcoxon rank-sum test). Right: N = 8 LVC mice, 9 V1 mice; R = −0.092; p = 0.724 (Pearson’s correlation test). **(D)** Top, posterior, and side views of LVCs (left) and V1 (right) lesions in two brains digitally reconstructed from laser-profilometry data. **(E)** Left: Same as in (C) but for the volumes of LVCs and V1 lesions measured from laser-profilometry reconstructions. N = 5 LVC mice, 5 V1 mice; *, p = 0.028 (Wilcoxon rank-sum test). Right: Scatter plot correlating V1 and LVC lesions areas with the Remapping Index. Symbols highlighted in green correspond to the examples in (D). N = 5 LVC mice, 5 V1 mice; R = −0.165; p = 0.790 (Pearson’s correlation test).

**Fig. S4.**
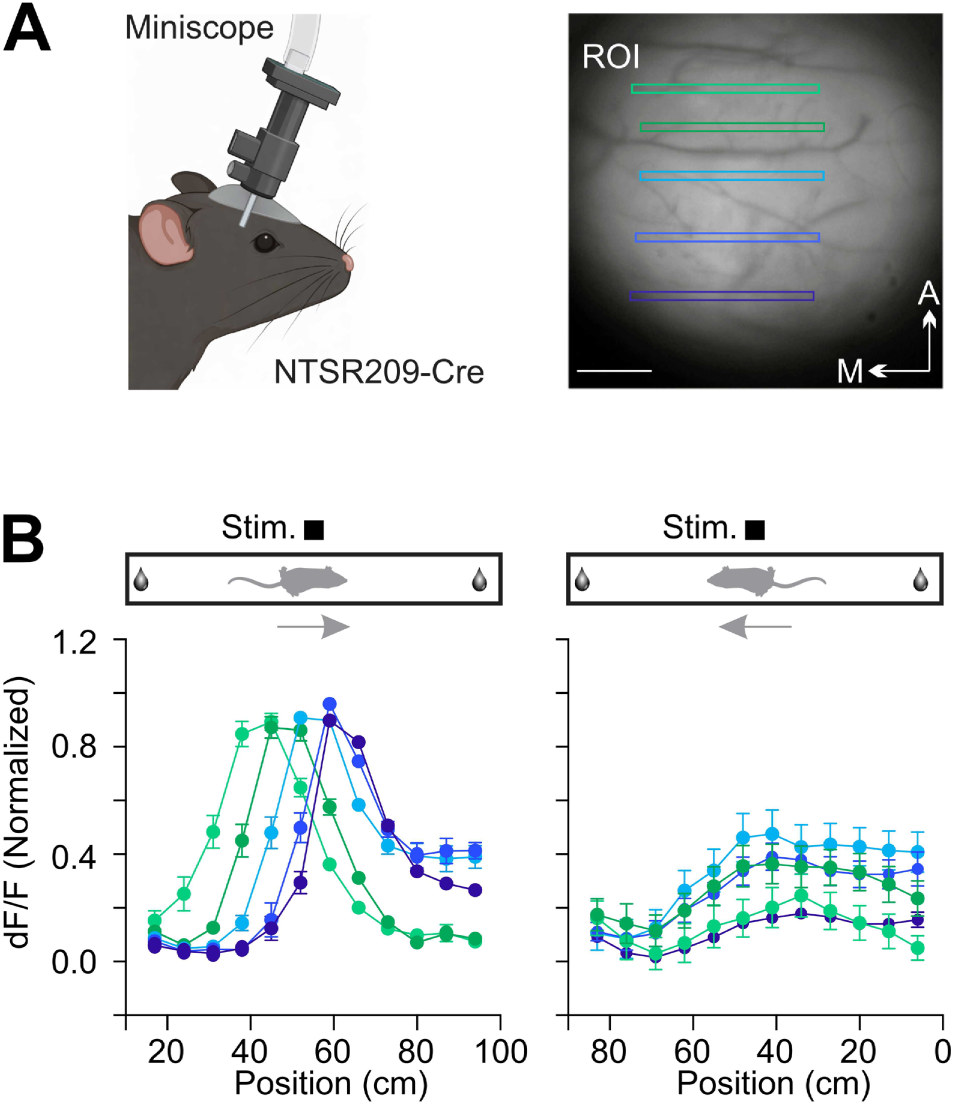
Hemifield specificity of SC widefield neurons during locomotion. **(A)** Left: Schematic of the experimental setup showing a Miniscope attached to the head of an NTSR1-GN209-Cre mouse. Right: Representative field of view with color-coded Regions of Interest (ROIs) drawn along the anterior-posterior axis of the SC. Scale bar: 200μm. **(B)** Top: Schematics of the linear track showing the position of a static visual stimulus (black square) while the mouse traverses the track in opposite directions. Bottom Left: Normalized fluorescence changes (dF/F) for color-coded ROIs plotted against the animal’s position on the track in the forward direction. Bottom Right: Normalized dF/F for the same ROIs plotted against position during traversal in the reverse direction. Error bars indicate SEM.

**Movie S1. Calcium imaging of the neuropil of widefield neurons in the superior colliculus during locomotion.**

Representative Miniscope imaging of the neuropil of the right superior colliculus (SC) of a Ntsr1–GN209–Cre mouse in which widefield neurons conditionally express GCaMP8m. The animal is traversing a linear track with a static visual cue (black square) positioned in the left hemifield. Note the spatial shift of the calcium transients as the animal approaches and passes the cue under illuminated conditions. These responses are absent in the dark. Thus, widefield cells respond to static stimuli moving across the visual field due to the animal’s locomotion. The video was recorded at 10 fps and is played back at 20 fps. Field of view: 1mm × 1mm.

